# Physiological and cognitive consequences of a daily 26h photoperiod in a primate *(M. murinus)*

**DOI:** 10.1101/2020.02.19.955674

**Authors:** Clara Hozer, Fabien Pifferi

## Abstract

Daily resetting of the circadian clock to the 24h natural photoperiod might induce marginal costs that would accumulate over time and forward affect fitness. It was proposed as the circadian resonance theory by Pittendrigh in 1972. For the first time, we aimed to evaluate these physiological and cognitive costs that would partially explain the mechanisms of the circadian resonance hypothesis. We evaluated the potential costs of imposing a 26h photoperiodic regimen compared to the classical 24h entrainment measuring several physiological and cognitive parameters (body temperature, energetic expenditure, oxidative stress, cognitive performances). We found significant higher resting body temperature and energy expenditure and lower cognitive performances when the photoperiodic cycle length was 26h. Together these results suggest that a great deviation of external cycles from 24h leads to daily greater synchronization costs, and lower cognitive capacities. To our knowledge, this study is the first to highlight potential mechanisms of circadian resonance theory.

## Introduction

Circadian rhythms, provide notable benefits to the organism compared to passive oscillations in local and transient response to exogenous factors (Sharma, 2003; Vaze and Sharma, 2013). The ubiquity of biological clocks in living organisms suggests a high adaptive value enhanced by the synchronization of behavioral and physiological processes with optimal phases of the cyclic environment (Sharma, 2003) and by the coordination of internal rhythms with each other (Pittendrigh, 1993; Paranjpe et al., 2003). The master clock controls vital metabolic cycles (Dvornyk et al., 2003) and synchronizes every day with the environmental *zeitgeber* (from german *zeit*: time and *geber*: giver), the most important being the light-dark cycles, whose period T is 24h. Light information is transmitted through retinal photoreceptors to the suprachiasmatic nuclei, where the central clock is based, which synchronizes the whole organism via chemical pathways, such as hormones (Moore, 2013; Buijs et al., 2013; Hankins et al., 2008; Yamaguchi et al., 2003).

Without any light clue, the circadian clock expresses its own periodicity, close to 24h, called the free-running period or *tau* (Halberg, 1962; Pittendrigh, 1960). *Tau* is both individual and species dependent (Pittendrigh and Daan, 1976) and maintained intra-cellularly by transcriptional and translational feedback loops regulating the expression of the clock genes (Duong et al., 2011; Lande-Diner et al., 2013). First proposed by Pittendrigh & Minis (1972), the circadian resonance theory assumes a relationship between *tau* and longevity: fitness is enhanced when the free-running period is close to the period of environmental cycles, *i.e.* when *tau* and T “resonate”. Indeed, drosophila reared under photoperiodic regimens far from 24h displayed reduced lifespan (Pittendrigh and Minis, 1972; von Saint Paul and Aschoff, 1978). These historical experiments were the first to confirm a negative link between the deviation of *tau* from 24h and longevity. Even though it seems very intuitive, the circadian resonance adaptive advantage is little shown in mammals. Wyse et al. (2010) found a negative correlation between the deviation of *tau* from 24h and longevity in several strains of laboratory mice, and several species of rodents and primates; Libert et al. (2012) showed that mice with circadian period close to 24h lived about 20% longer than those with shorter or longer *tau.*

These two studies provide evidence that keeping a 24h free-running period positively affects lifespan. However, the underlying mechanisms that would explain the disadvantages of a desynchrony between *tau* and external light-dark cycles remain totally unknown. An assumption advanced is that daily marginal metabolic or physiological costs, required by the clock daily entrainment, would accumulate over life and with time, impact negatively longevity, according to the rate of living theory. Postulated by Pearl (1928), this theory states that longevity of an organism is conditioned by its rate of metabolism. Our study aimed at determining if these daily costs could be detected and quantified.

To address this issue, we focused on a non-human primate, the gray mouse lemur (*Microcebus murinus*). This small Malagasy lemur is an emerging model in neurosciences since it displays aged-related impairments similar to those found in humans (e.g. spontaneous neurodegenerative diseases or cognitive deficiencies, Bons et al., 2006; Joly et al., 2014; Languille et al., 2012; Picq et al., 2015), including circadian rhythms alteration, such as locomotor activity fragmentation or sleep deterioration (see Hozer et al., 2019 for review). On the other hand, its small body size and light body mass make it an ideal and promising laboratory model. Regarding circadian features, the gray mouse lemur is strictly nocturnal and its metabolism is highly dependent on photoperiod (Génin and Perret, 2000). Its mean free-running period lies around 23.5h (Cayetanot et al., 2005).

We submitted mouse lemurs to two different photoperiodic regimens, mimicking a standard deviation of *tau* (individuals kept in light-dark cycles of 24h) and a great deviation of *tau* (individuals kept in light-dark cycles of 26h). We considered that clock daily entrainment may exert a direct or indirect influence on physiological and metabolic functions, particularly on basal metabolic activity, which might affect longevity. We thus focused on several factors that reflect this basal metabolism (e.g. oxygen respiration, energy expenditure, body temperature, oxidative stress…) and measured them before and after photoperiodic treatments. Knowing that the circadian clock substantially influences cognitive performances (Kyriacou and Hastings, 2010), we were also wondering if daily synchronization costs could affect cerebral abilities. We hence assessed cognitive performances using a learning-task based on visual discrimination.

## Results

### 1) Baseline before treatment

Baseline parameters are shown in Table 1. No difference was detected between the two groups before treatments. All values (except for oxidative stress) are averaged over the five first days before treatment.

**Table 1:** Body mass, oxidative stress and energy parameters, body temperature and locomotor activity during resting and active phases. a.u. = arbitrary unit.

| Parameters | T24 (n=11) | T26 (n=11) | F | P |
| --- | --- | --- | --- | --- |
| Body mass (g) | 85.1±1.7 | 84.0±2.3 | 0.02 | 0.88 |
| Oxidative stress (ng.mL<sup>-1</sup>) | 49.34±9.67 | 57.33±9.72 | 0.54 | 0.47 |
| **Resting phase** |  |  |  |  |
| VO<sub>2</sub> (mL.kg<sup>-1</sup>.h<sup>-1</sup>) | 1599.9±63.5 | 1636.0±67.0 | 0.24 | 0.63 |
| VCO<sub>2</sub> (mL.kg<sup>-1</sup>.h<sup>-1</sup>) | 1328.9±49.6 | 1304.3±51.2 | 0.24 | 0.63 |
| Heat (kcal.h<sup>-1</sup>) | 0.66±0.03 | 0.69±0.03 | 0.91 | 0.35 |
| Body temperature (°C) | 36.40±0.24 | 36.38±0.27 | 0.01 | 0.93 |
| Locomotor activity (a.u.) | 1.66±0.17 | 2.64±0.42 | 1.39 | 0.26 |
| **Active phase** |  |  |  |  |
| VO<sub>2</sub> (mL.kg<sup>-1</sup>.h<sup>-1</sup>) | 2627.2±214.6 | 2737.2±164.4 | 0.01 | 0.95 |
| VCO<sub>2</sub> (mL.kg<sup>-1</sup>.h<sup>-1</sup>) | 2576.1±200.4 | 2611.2±151.1 | 0.18 | 0.68 |
| Heat (kcal.h<sup>-1</sup>) | 1.13±0.07 | 1.17±0.08 | 0.01 | 0.94 |
| Body temperature (°C) | 38.05±0.14 | 37.86±0.16 | 0.67 | 0.43 |
| Locomotor activity (a.u.) | 20.46±5.50 | 18.92±3.18 | 0.19 | 0.67 |

### 2) Effects of photoperiodic treatments on locomotor activity and body temperature

T24 and T26 animals responded properly to their respective photoperiodic treatments (locomotor activity mean periods of 23.99±0.01 and 25.99±0.04 respectively), since they were correctly entrained to the light-dark cycles they were submitted to (t_T24_=-2.53, p_T24_=0.35, t_T26_=-0.75, p_T26_=0.47; Fig. 1). The diagrams of temperature (see Supplementary materials S1) present the same patterns (mean periods of 24.01±0.01 and 25.98±0.07 respectively; t_T24_=1.83, p_T24_=0.15; t_T26_=-0.80, p_T26_=0.44).

**Figure 1:**
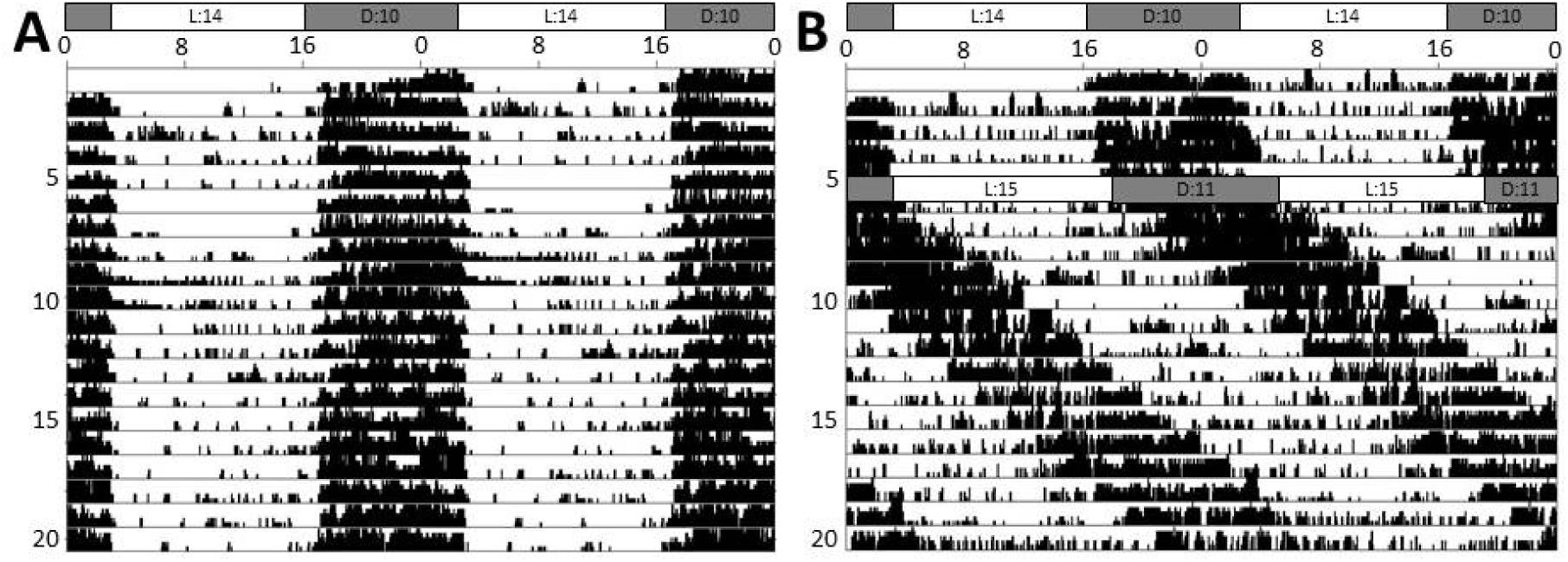
Representative double-plot actograms of individuals of each photoperiodic treatment during the 20 first days of experiment. Before treatment, all individuals were living under light-dark cycles of 24h with 14h of light and 10h of dark (L:D 14:10). The different treatments started on the 6th day of recording. A: T24 group. Mouse lemurs were kept under L:D 14:10. B: T26 group. Mouse lemurs were kept under light-dark cycles of 26h (L:D 15:11).

No significant difference was observed in activity levels between T24 and T26 animals (active phase: F=1.08, p=0.32; resting phase: F=0.04, p=0.85; Fig. 2). Regarding body temperature, a rising trend in T26 was detected only during the resting phase (active phase: F=2.08, p=0.17; resting phase: F=2.99, p=0.11; Fig. 2). When subtracting the thermic fall, *i.e.* considering only Tb values starting after the daily minimal Tb (T_min_) was reached, T26 animals exhibited significantly higher Tb than T24 (+0.32±0.06°C, F=8.63, p=0.009, Fig. 2). T_min_ was reached 3.33h and 3.20h after the start of the resting phase in T24 and T26 animals respectively.

**Figure 2:**
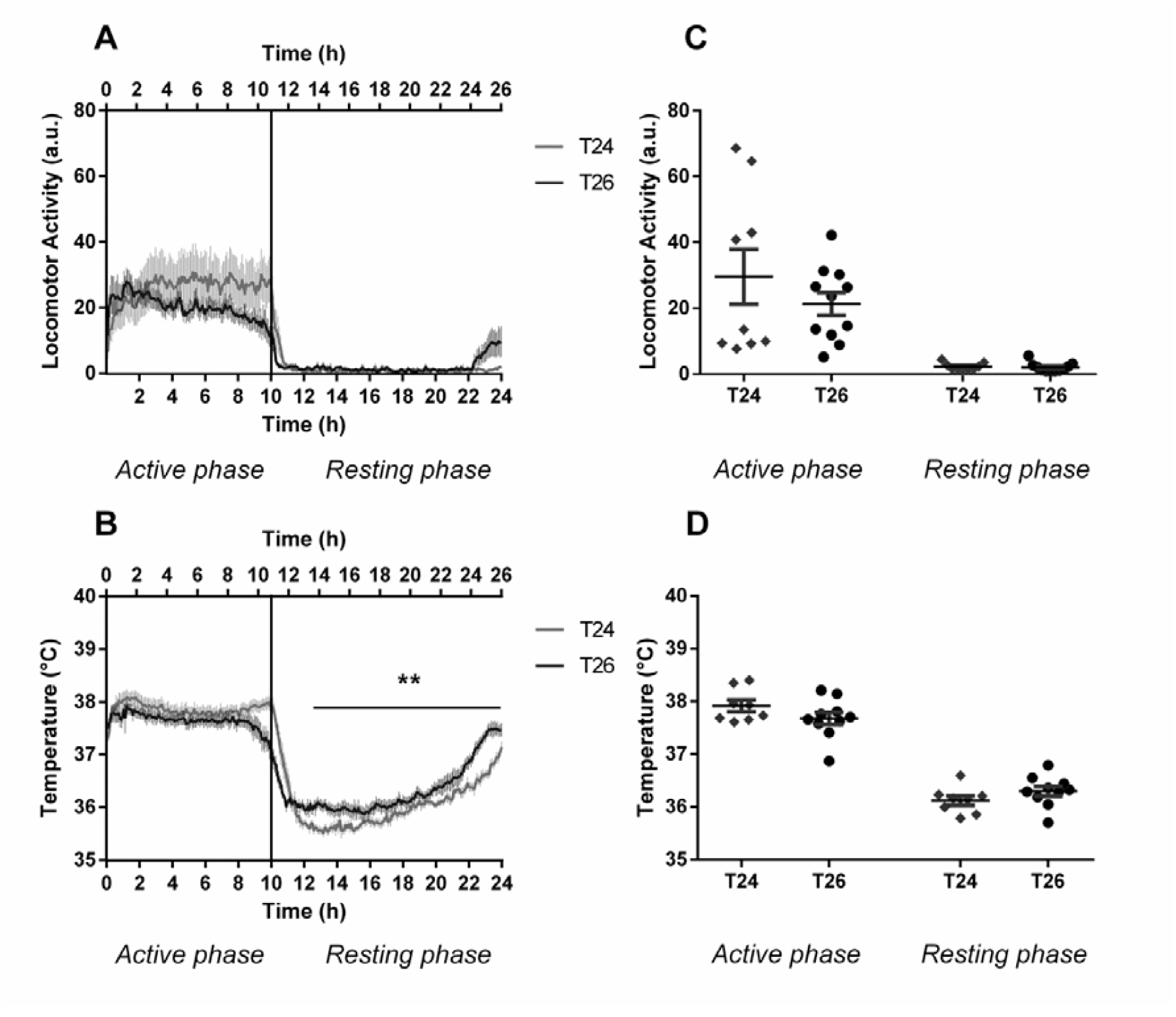
A & B: Daily profiles of locomotor activity and body temperature in T24 and T26. Switch between night and day is represented by the black line. Graphs of T26 animals were scaled to fit the T24 animals’ ones. No significant difference in activity levels was observed between two treatments. T26 animals had significant higher Tb during resting time compared to control animals T24 after the thermic fall. C & D: Mean locomotor activity and body temperature during active and resting periods in T24 and T26. ‘**’=p<0.01

### 3) Effects of photoperiodic treatments on calorimetric parameters

When considering the active phase, VO_2_, VCO_2_ and Heat levels were not significantly different between treatments (F=0.33, p=0.58; F=0.06, p=0.82; F=0.62, p=0.44 respectively; Fig. 3). During the resting phase, there is a tendency to higher oxygen consumption (F=3.77, p=0.07), confirmed by significantly higher VCO_2_ (+289.3±107.4 ml.kg^-1^ .hr^-1^, F=8.83, p=0.009) and Heat (+0.10±0.06 kcal.h^-1^, F=5.16, p=0.04) in T26 compared to T24 (Fig. 3). Both groups did not exhibit any significantly different body masses at the end of the treatments (F=0.15, p=0.70).

**Figure 3:**
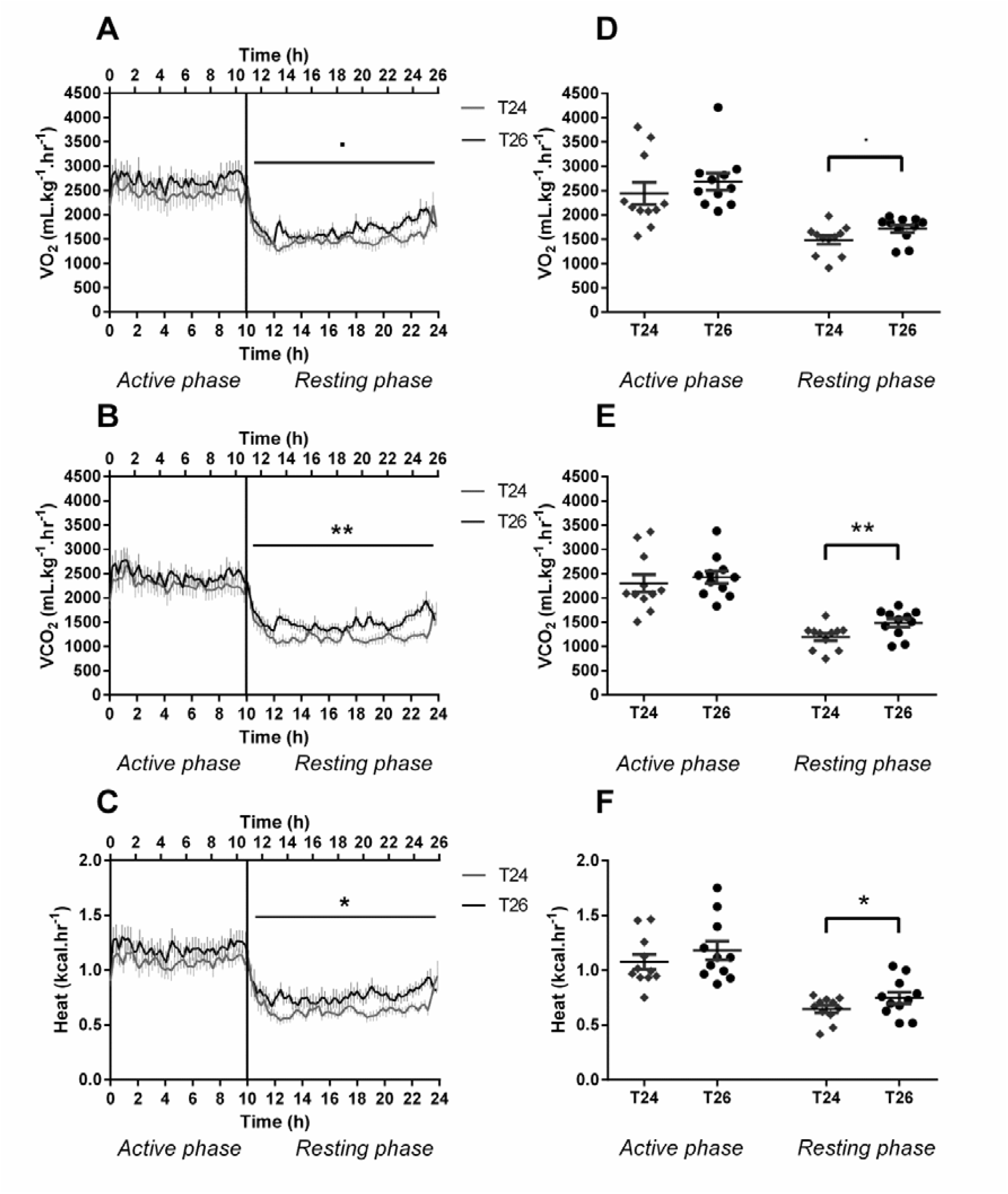
A, B & C: Daily profiles of VO_2_, VCO_2_ and Heat in T24 and T26. Switch between night and day is represented by the black line. Graphs of T26 animals were scaled to fit the T24 animals’ ones. D, E & F: Mean VO_2_, VCO_2_ and Heat during active and resting periods in T24 and T26. A tendency suggests a higher VO_2_ during the resting phase in T26 and significantly higher VCO_2_ and Heat during the resting time were observed in T26. ‘.’=p<0.1, ‘*’=p<0.05, ‘**’=p<0.01.

### 4) Oxidative stress

Plasma 8OHdG levels were not significantly different between groups (47.71±20.21 vs 51.30±22.75 ng.mL^-1^, F=0.13, p=0.72, Fig. 4).

**Figure 4:**
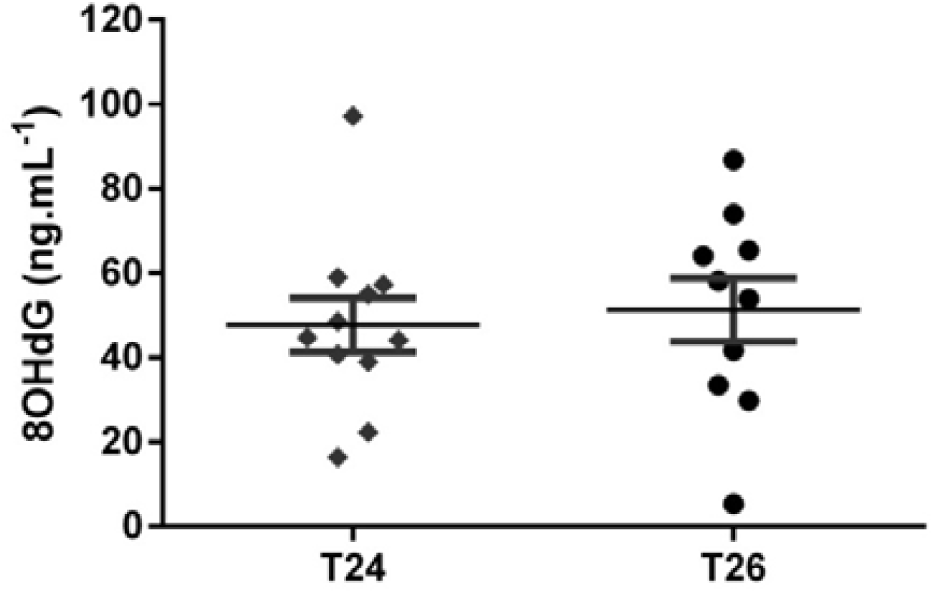
Plasma 8OHdG levels in T24 and T26 (ng.mL^-1^). No significant difference was observed between both treatments.

### 5) Learning performances

Among the 22 animals, 7 reached the success criterion: 3 in T24 and 4 in T26. The T26 individuals needed significantly more trials to reach the criterion than the T24 (17.5±1.5 vs 7±3.33, p=0.033; Fig. 5).

**Figure 5:**
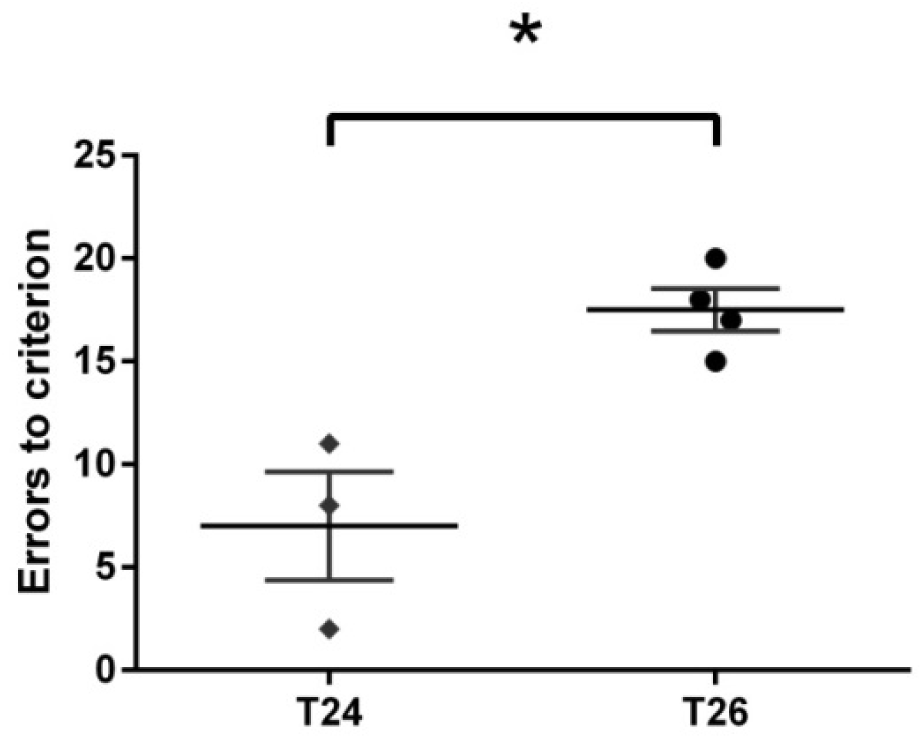
Trials needed before reaching the success criterion to the learning task. T26 individuals needed significantly more trials than T24 individuals to reach the criterion. ‘*’=p<0.05

### 6) Correlations between physiological, metabolic and cognitive parameters

In addition to the fact that most of calorimetric parameters are strongly correlated, oxidative stress is also positively correlated and tends to be correlated with the resting VO_2_ and VCO_2_ respectively. Furthermore, the number of trials needed to reach the success criterion in the cognitive task is positively correlated to resting VO_2_ and VCO_2_, *i.e.* the higher resting VO_2_ and VCO_2_, the worse the cognitive capacities are. Finally, body temperature after minimal temperature was reached was positively correlated with resting VCO_2_, and tends to correlate positively with resting VO_2_ and negatively with cognitive performances (Table S1).

## Discussion

In the present study, we aimed at investigating the metabolic costs of the circadian clock daily synchronization and its cognitive consequences. Our results suggest that a physiological, metabolic and cognitive cost is daily generated when the free-running period *tau* does not match the environment periodicity of 24h. Animals submitted to light-dark cycles of 26h, mimicking a great divergence between *tau* and external cycles, exhibited higher resting body temperature, higher VO_2_, VCO_2_ and Heat levels and lower cognitive performances. To our knowledge, this is the first time that a study provides some insights on the potential biological costs of clock daily synchronization. The potential costs of synchronization had been first evoked by Pittendrigh and Minis (1972), then by Wyse et al. (2010), in order to elucidate the circadian resonance mechanisms, but had never been assessed so far.

Animals in T24 and T26 responded correctly to their light-dark treatments and random feeding times over the day prevented the animals from being entrained by food distribution. Nevertheless, we verified that T26 animals were properly entrained to their photoperiodic regimen without masking effect. Although locomotor activity was restricted to daily dark period, it cannot be ruled out that other internal body rhythms may be desynchronized with external environment. Tb rhythm of T26 individuals displayed a period of 26h, in other words, was entrained to the light-dark cycles and did not free-run. Despite its non-independent link with locomotor activity, body temperature can inform on organism entrainment. In most experiments where T is modified and individuals’ rhythms decoupled from external cycles, body temperature and locomotor activity are not synchronized, locomotor activity being restricted to activity periods and body temperature free-running (Hasan et al., 2018; Karatsoreos et al., 2011; Legates et al., 2012; Leise et al., 2018; Vivanco et al., 2010), which was not the case in our data. Moreover, experiments on entrainment limits in mouse lemurs conducted in our laboratory show that locomotor activity and temperature start decoupling over 27h (M. Perret, personal communication, dec. 2019), suggesting that 26h T-cycles lie within the mouse lemur range of entrainment.

We observed that T26 animals displayed significantly higher body temperature and higher energy expenditure (VO_2_, VCO_2_ and Heat) during the resting phase, especially after the daily metabolic fall. Higher body temperature was neither the consequence of a higher locomotor activity during the same period. Nor is it due to a potential adaptation time to 26h cycles because the effect was found regardless of the considered week, even after 3 weeks of photoperiodic treatment (data not shown). A lot of studies on modified T-cycles were carried out on various species but only a few of them dealt with daily metabolic and physiological consequences of the organism’s entrainment to T-cycles different from 24h, and none focused on metabolic rate or body temperature. In invertebrates, two species of *Camponotus* ants exhibited significantly faster pre-adult development under T24 compared toT20 and T28, suggesting a fitness advantage of “resonating” clock (Lone et al., 2010). Short-period mutant hamsters (whose “natural” endogenous period lies around 24.05h, Davis and Viswanathan, 1992) displaying a 22h free-running period died younger and exhibited severe cardiac and renal diseases when raised under cycles of 24h (Martino et al., 2008). Vilaplana et al. (1995) showed that rats kept under T25 and T26 light-dark cycles during 64 days exhibited lower body weight, less food intake and less efficiency (*i.e.* relationship between body weight increase and food intake) than rats kept under T24 cycles. This last experiment could draw an interesting parallel with our results, since assessed parameters are comparable to those we measured. Nevertheless, the theory behind these observations is actually totally opposite to our own hypothesis. Indeed, Vilaplana and colleagues started from the postulate that cycles of 25 or 26h are closer from the rat’s free-running period that 24h and that rats better “resonate” under this cycle length. In this regard, higher available energy is preferentially allocated to activity and metabolism, rather than to growth efficiency. On the contrary, we suggest that individuals kept close from resonance frequency would consume less energy to maintain their clock reset and display a lower metabolism, which contradicts Vilaplana’s hypothesis. Furthermore, the hypothesis of a “better resonating clock under T25 and T26 cycles” can be questioned, since the free-running period of rat is approximately 24.5h (Jan Stenvers et al., 2016; Strijkstra et al., 1999; van Gool et al., 1987).

Among endotherms it has been widely accepted that body temperature is directly influenced by metabolic rate, as this is the principal origin of endogenous heat (Berger et al., 1988; Clarke et al., 2010; Gillooly et al., 2001; Lovegrove, 2003; Rey et al., 2015; Rezende and Bacigalupe, 2015; Rising et al., 1992). In this respect, our calorimetric results are in line with body temperature observations. Indeed, VO_2_, VCO_2_, Heat and Tb were significantly higher in T26 than in T24 animals during the resting time that closest resembles the basal metabolic state of the animal. Intraspecific negative relationships between longevity, aging and basal metabolic rate have been found out in numerous species. They corroborate the rate of living theory (Conti et al., 2006; Redman et al., 2018; Van Raamsdonk et al., 2010), although comparisons across species sometimes show that metabolism is not correlated to lifespan (Niitepõld and Hanski, 2013; Promislow and Haselkorn, 2002; Robert et al., 2007), especially in birds and bats (Munshi-South and Wilkinson, 2010). As for body temperature, the relationship with aging is well known and fully documented. Several studies confirm that lower body temperature leads to slower aging and longer lifespan in poikilotherms as well as in homeotherms (Conti, 2008). Hcrt-UCP2 mutant mice, displaying a reduction of Tb of 0.5 – 0.6 °C demonstrated up to a 20% increase of median lifespan (Conti et al., 2006). Male C57B1/6 mice tended to live longer than females and displayed a significantly lower body temperature of 0.2 to 0.5 °C (Sanchez-Alavez et al., 2011). In humans, the Baltimore Longitudinal Study of Aging (BLSA) reported a lower body temperature related to higher lifespan and other positive physiological effects (Shock et al., 1984).

The rate of living theory has then been further extended into the free-radical theory of aging stating that aging results from accumulating damages produced by reactive oxygen species, such as 8OHdG (Harman, 1956). Our results did not exhibit any significant divergent plasma 8OHdG levels between the different photoperiodic groups. The light-dark treatments provided to the animals may have not been compelling or long enough to impact significantly the individuals at the DNA level. It highlights that daily costs imposed by the circadian clock resetting remain marginal at the cellular level. Further investigations could then consist in extending the duration of the metabolic stress imposed by the light-dark cycles.

Regarding cognitive outcomes, only seven animals reached the success criterion to the learning task, which is one-third of the total number of individuals. This result may seem low but it lies close to the success rate observed in other experiments using the same cognitive apparatus (Picq et al., 2015; Royo et al., 2018). The resulting interpretations must though be viewed cautiously. Among animals that learned the task, T26 animals needed significantly more trials to reach the success criterion than the T24. A higher required energy to reset the circadian clock may create a side-effect in cognitive performances, highlighting a potential trade-off between metabolism and cerebral abilities. It is well documented that a strong link exists between circadian clock and cerebral performances. Through its link with the hippocampus (Borgs et al., 2009; Chiang et al., 2017; Schnell et al., 2014, Snider et al., 2018), the circadian system influences mood, learning, time-place association and memory in laboratory mice (Albrecht, 2017; Cain et al., 2004; Legates et al., 2012; Ruby et al., 2008), and the involvement of clock genes is well established (Snider et al., 2016; Van der Zee et al., 2008; Wang et al., 2009). Furthermore, numerous data indicate that circadian disorganization (jet-lag, phase shifts, aging alterations, sleep impairments, shift work…) invariably leads to impaired cognitive performances which suggests that clock resynchronization indirectly impacts cognitive capacities (Antoniadis et al., 2000; Chellappa et al., 2018; Gibson et al., 2010; Krishnan and Lyons, 2015; Loh et al., 2010; Rouch et al., 2005). Only a very few studies have tested the effects of a chronic misalignment between *tau* and T, without sleep-wake cycles decoupled from circadian rhythm. Neto et al. (2008) reported decreased performances in a passive avoidance memory task in rats kept under 22h light-dark cycles compared to control group (L:D 24h). The authors of this study raised the involvement of the circadian clock in an emotional component of the memory task, related to fear or risk evaluation. The cognitive cost observed in T26 animals could also suggest a lack of sleep, or at least sleep modification. Locomotor activity profiles are not significantly different between T26 and T24 groups even though there seems to be a period between 24h and 26h during which T26 animals activate (Fig. 2A). This non resting period could be associated with a slight sleep debt, which, cumulated over 20 days, could alter learning performances. A prolonged sleep deprivation affects cognitive performances, as it has been shown in humans (Reynolds and Banks, 2010; K. J. Wright et al., 2006), even if the sleep deprivation is moderate (Sadeh et al., 2003; Van Dongen et al., 2003). In this perspective, a cumulative sleep debt due to sooner activation of T26 animals could explain the worse results during the discrimination task, without being directly related to a daily synchronization cost. The altered cognitive performances of T26 animals may also be due to an improper entrainment to the imposed light-dark cycles. Although locomotor activity and body temperature were synchronized with each other, other internal rhythms might be desynchronized, suggesting a masking effect of light on locomotor activity and correlated body temperature, as previously mentioned. A lot of studies report the negative effect of body rhythms desynchrony on neuro-behavioral functions, especially during aging, when the biological clock undergoes severe alterations (Cajochen et al., 2004; Colwell, 2015; Grady et al., 2010; Krishnan and Lyons, 2015; Wright et al., 2006b). In that case, weaker learning performances in T26 animals would rather be due to a potential internal desynchrony rather than to a cost of synchronization.

Although the inter-correlations between cognitive, cellular and metabolic parameters suggest a multi-scale effect of photoperiodic treatments on tested individuals, the precise mechanisms underlying the relationship between clock daily synchronization and metabolic costs remain so far unknown and can only be hypothesized. They besides require the cellular mechanisms of light entrainment, that remain barely investigated (Johnson, Elliot & Foster, 2003). In laboratory conditions, *i.e.* under constant intensity of light, clock entrainment is supposed to follow a discrete model, where light activation and extinction are supposed to mimic dawn and dusk transitions. The light is indeed assumed to act effectively at phases transition, e.g. at dawn or dusk. In the case of the T26 animals, synchronization of clock should occur at the beginning of the light phase (*i.e.* after 11h of darkness) and at its end (*i.e.* after 15h of light). As described by Johnson, Elliot & Foster (2003), the entrainment of the clock should induce a great phase delay at dusk (*i.e.* light extinction) and a smaller phase advance at dawn (*i.e.* light activation), in such a way that the net phase shift is (26h - *tau*), that is around 2h. In that configuration, the light input is postulated to activate the expression of Fos genes, followed by the activation of Per1 and Per2, dependently on the moment of the circadian time (Cao et al., 2015; Challet, 2007). It has also been suggested that epigenetic processes were involved in clock synchronization and plasticity (Azzi et al., 2017). These reactions are energy-consuming (Bass and Takahashi, 2010; Golombek and Rosenstein, 2010), and one may suppose that the energy required to elicit an important phase delay in the T26 animals is greater than the energy required for the resynchronization of the T24 ones. High energetic gene activations may lead to higher metabolism and as a consequence, higher body temperature. A further interesting question is to know whether a continuous model of entrainment (*i.e.* with transitions between light and dark phases and daily varying illumination levels) would affect metabolic features differently.

To conclude, our results seem to highlight that external photoperiod lengths deviating from the free-running period increase metabolic daily requirements to keep the circadian clock reset and might result in a cognitive cost. The accumulation of repetitive marginal costs would accelerate aging process and lead to lower survival. These investigations could then provide an initial insight of the mechanisms underlying the circadian resonance theory and opens the way to further investigations: immune, cardiovascular or other markers of fitness, such as reproduction performances could constitute complementary significant costs of circadian clock daily synchronization.

## Materials & Methods

### 1) Animals and ethical statement

All mouse lemurs studied were males born in the laboratory breeding colony of the CNRS/MNHN in Brunoy, France (UMR 7179 CNRS/MNHN; European Institutions Agreement # E91–114.1). All experiments were performed in accordance with the Principles of Laboratory Animal Care (National Institutes of Health publication 86-23, revised 1985) and the European Communities Council Directive (86/609/EEC). The research was conducted under the approval of the Cuvier Ethical Committee (Committee number 68 of the “Comité National de Réflexion Ethique sur l’Expérimentation Animale”) under authorization number 12992-2018011613568518 v4.

All cages were equipped with wood branches for climbing activities as well as wooden sleeping boxes. The ambient temperature and the humidity of the rooms were maintained at 25 to 27°C and at 55% to 65%, respectively. When they are not involved in experimental protocols, animals in facilities are exposed to an artificial photoperiodic regimen consisting of alternating periods of 6 months of summer-like long day (Light-Dark (L:D) 14:10) and 6 months of winter-like short days (L:D 10:14), in order to ensure seasonal biological rhythms (Perret and Aujard, 2001). The following described experiment focused on animals during the long-day season.

### 2) Experimental procedure

Twenty-two male gray mouse lemurs between 2 and 4 years-old were chosen randomly in the colony. Before the start of the experiment (see Fig. 6), the animals were isolated under light-dark cycles of 24h (L:D 14:10); the metabolic activity was measured using a calorimetry system during 5 days, followed by blood sampling to assess individual oxidative stress. We then submitted the animals to two different photoperiodic treatments during 22 days. Individuals of the first group (T24, n=11) were kept under light-dark cycles T of 24h with 14 hours of light (resting phase) and 10 hours of darkness (active phase) per day (L:D 14:10). Mouse lemurs of the second group (T26, n=11) were submitted to light-dark cycles T of 26h (L:D 15:11), mimicking a great divergence between their free-running period and the environmental periodicity. After 15 days, metabolic activity was measured again during 5 days in the indirect calorimetry set-up, followed by blood sampling. Afterwards, the animals performed a cognitive task during the last 2 days of experiment. Animals were fed at random times of the day with fresh fruits and a homemade mixture, corresponding to an energy intake of 25.32 kcal.day^-1^ (see Dal-pan et al., 2011 for details). Food intake and body mass were daily monitored, and body temperature (Tb) and locomotor activity (LA) were recorded continuously using telemetry implants during the whole experiment.

**Figure 6:**
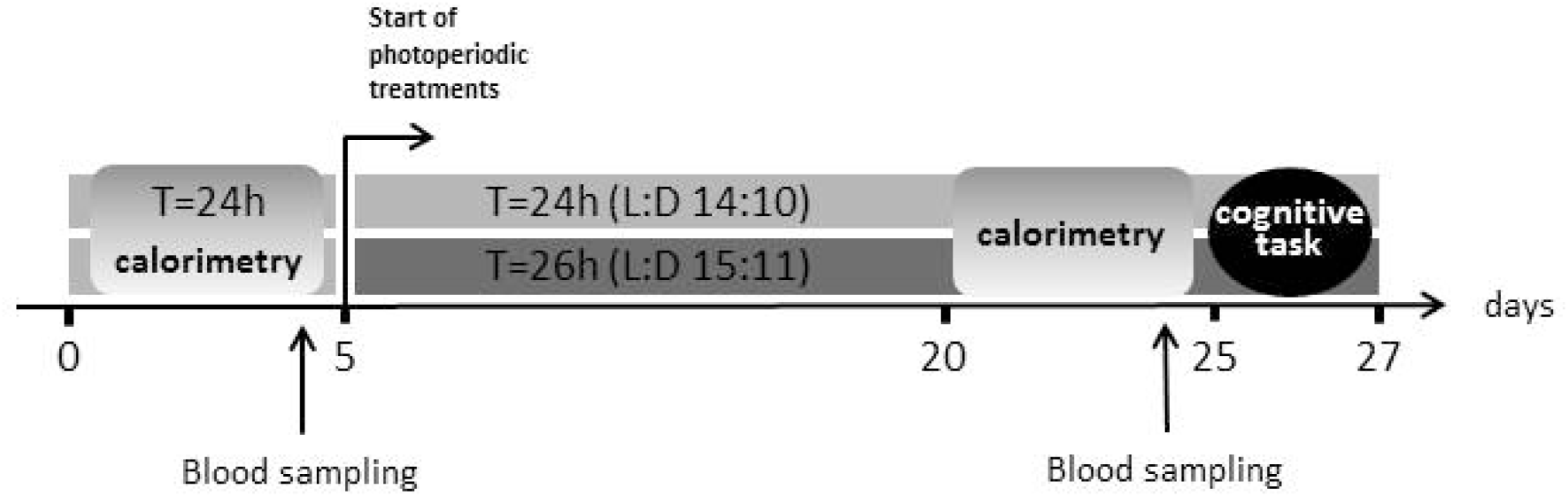
Experimental design. Animals’ metabolic activity (calorimetry) was first assessed before treatments during 5 days, period during which pre-treatment blood was also collected. The animals were then separated into two different photoperiodic regimens: in light-dark cycles of 24h (L:D = 14:10) or in light-dark cycles of 26h (L:D = 15:11). After 15 days, metabolic activity was measured again and new blood sampling was collected. The animals then performed a learning task during 2 days.

### 3) Indirect calorimetry system

Metabolic activity was recorded using an indirect calorimetry system (Oxymax, Colombus Instruments Inc, Columbus, Ohio, USA). Animals were housed in individual metabolic cages during 5 days (with one day for acclimation before measurements). Oxygen consumption (VO_2_), carbon dioxide production (VCO_2_) and energy expenditure (*Heat* = 3.815 × *VO*_2_ + 1.232 × *VCO*_2_, see Lusk, 1924) were recorded continuously. VO_2_ and VCO_2_ were expressed as a function of the whole body mass (mL.hr^-1^ .kg^-1^) and Heat in kcal.hr^-1^.

### 4) Body temperature and locomotor activity telemetric monitoring

Recording of locomotor activity and body temperature was obtained by telemetry. A small telemetric transmitter weighing 2.5 g (model TA10TA-F10, DataScience Co. Ltd, Minnesota, USA) was implanted into the visceral cavity under isoflurane anesthesia (4% for induction and 1-1.5% for maintenance). After surgery, animals returned to their home cage and were allowed to recover for 15 days before the start of experiment and continuous recordings of LA and Tb. Total recovery was checked by visual inspection of the complete healing of the surgical incision and by verification of a stable daily pattern of Tb variations. Temperature was punctually monitored every 5 minutes and locomotor activity was continuously recorded by a receiver placed into the cage, which detected vertical and horizontal movements and transmitted data to the computer (coordinate system, DataquestLab Prov.3.0, DataScience Co.Ltd, Minnesota, USA). LA data were summed in 5 minutes intervals and expressed in arbitrary unit (a.u.). Periods of activity rhythms were calculated using the Lomb–Scargle periodogram (LSP) procedure (Ruf, 1999), with Clocklab software (Actimetrics, Evanston, IL, USA). AL and Tb values were averaged over the 20 days of photoperiodic treatment (the last two days were excluded to avoid the influence of the cognitive task on AL and Tb data). Two implants in the T24 group and one implant in the T26 group were found partially or totally defective; Tb and AL could not thus be recorded.

### 5) Oxidative stress measurements

Mouse lemurs’ blood was collected 3h before the onset of the individuals. Two hundred µL of blood were taken via the saphenous vein, and collected in tubes containing EDTA. Blood samples were then centrifuged at 2000g at 4°C for 30 minutes and plasma was collected in order to measure plasma 8-hydroxy-2’-deoxyguanosine (8-OHdG) levels in ng.mL^-1^ (OxiSelect™ Oxidative DNA Damage Elisa kit, Cell Biolabs Inc.).

### 6) Cognitive apparatus

The cognitive task was first described by Picq et al. (2015), inspired from apparatus designed by Lashley (1930) for rodents and based on visual discrimination. In the present study, it was conducted over a two-day period, 3h hours before the light extinction. The first day is dedicated to animals’ habituation to the set-up. The discrimination test takes place on the second day. The animal is introduced in a big squared vertical cage through an opening in the wall, to an elevated starting platform. It has to jump onto one of two landing platforms, fixed below on the opposite wall. A hole centered behind the two landing platforms leads to a nesting box behind the cage. On each landing platform, a visual *stimulus* helps discriminating the left and right platforms. One of these visual clues is the positive *stimulus*, the other is the negative one. At each trial, the landing platform with the positive clue is kept fixed and leads to the nesting box, whereas the platform with the negative clue is mobile and toggles when the animal jumps on it, such a way that it falls on a cushion pillow, to prevent any injury. At each trial, the location of the positive and negative stimuli on the right or left landing platform is randomized. The animal was given 30 trials to reach the success criterion consisting in 8 jumps on the positive platform within 10 consecutive trials. For each animal, we measured the number of trials needed to reach the success criterion (for more details, see Picq et al., 2015).

### 7) Statistical analysis

Normality was verified using Shapiro-wilk tests. Non-normally distributed data were log-transformed (*i.e.* Heat of the active phase and locomotor activity). Outliers were detected and removed using Dixon tests (one T26 individual for oxidative stress). T-tests were used to verify the proper entrainment of T24 and T26 animals. ANOVAs between different treatments were performed for the mean VO_2_, VCO_2_, Heat, Tb and LA during active phase and resting phase, body mass and oxidative stress before and after treatments. A non-parametric Kruskal-Wallis test was applied to compare cognitive performances between the two treatments. Data were analyzed using R Studio software with p<0.05 taken as statistical significance. Data are presented as mean ± SEM.

## Acknowledgments

We warmly thank Jérémy Terrien for his assistance in the statistical analysis and his thorough re-reading of this article and Martine Perret and Fabienne Aujard for their help in elaborating the experimental design.

