## Supplementary materials for "Physiological and cognitive consequences of a daily 26h photoperiod in a primate *(M. murinus)*"

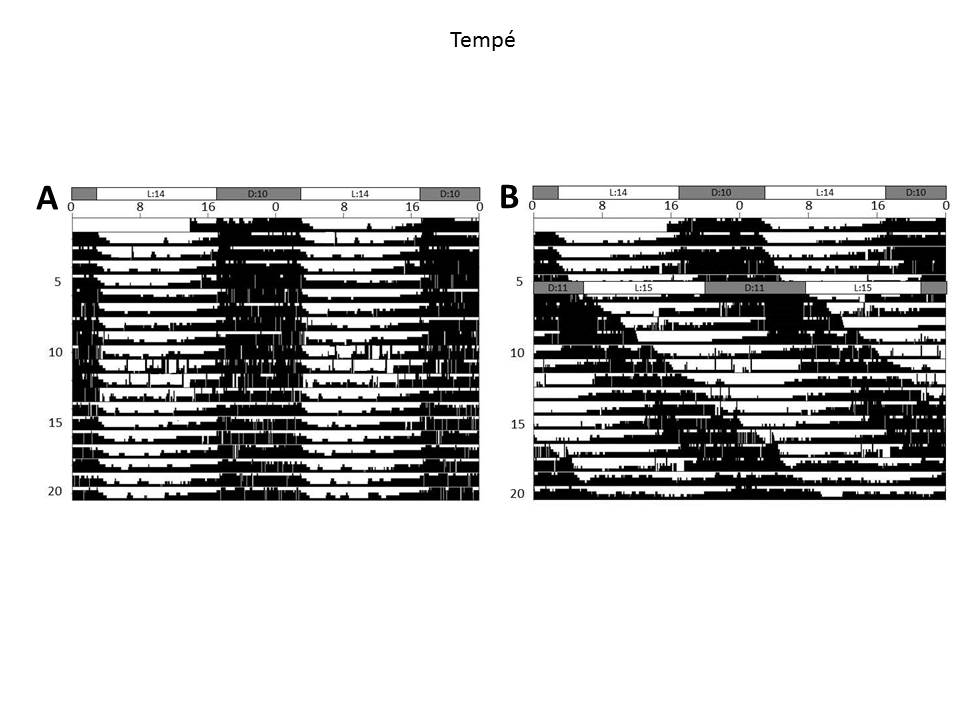
Figure S1: Representative double-plot temperature diagrams of individuals of each photoperiodic treatment during the first 20 days of experiment. Before treatment, all individuals were living under light-dark cycles of 24h with 14h of light and 10h of dark (L:D 14:10). The different treatments started on the 6th day of recording. A: T24 group. Mouse lemurs were kept under L:D 14:10. B: T26 group. Mouse lemurs were kept under light-dark cycles of 26h (L:D 15:11).

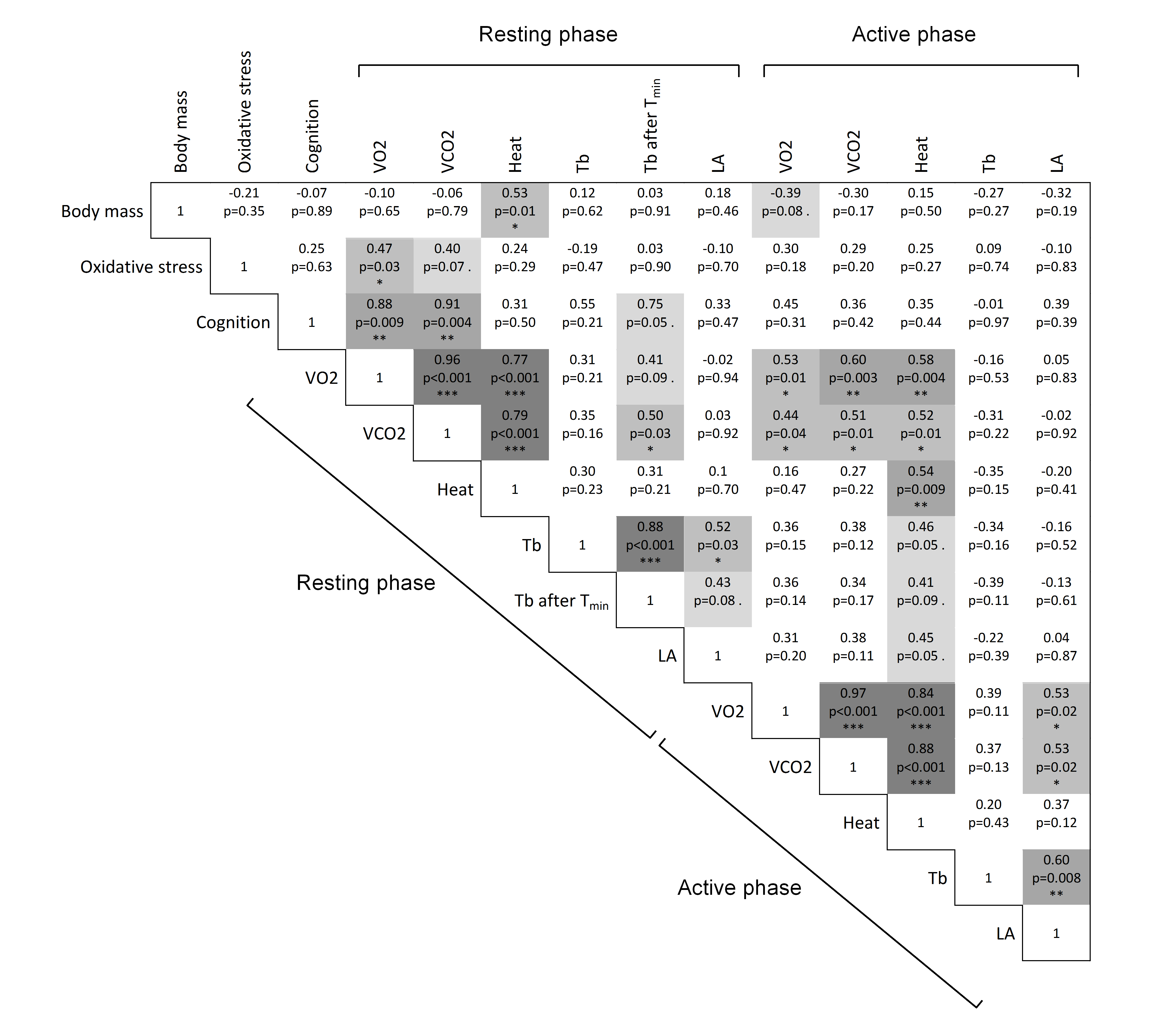

Table S1: Correlation table of all considered variables. Cells are colored depending on significance levels. Most of calorimetric parameters are strongly correlated. Oxidative stress is positively correlated with resting VO_2_ and tends to be correlated with resting VCO_2_. Cognitive capacities (*i.e.* number of error before criterion) are positively correlated with resting VO_2_ and VCO_2_ and tend to be correlated with Tb after T_min_. Tb after T_min_ is also positively correlated with resting VCO_2_ and tends to be positively correlated with resting VO_2_ and active Heat.
